# Developing Small Molecules that Inhibit K-Ras/GTP Binding Based on New Affinity Measurements

**DOI:** 10.1101/2020.07.27.218248

**Authors:** Luca Carta, Rebecca Hutcheson, Simon A. Davis, Michael J. Rudolph, Charles H. Reynolds, Matthias Quick, Theresa M. Williams, Michael Schmertzler, Yaron R. Hadari

## Abstract

*RAS* genes encode small GTPases essential for proliferation, differentiation, and survival of mammalian cells. *RAS* gene mutations are associated with approximately 30% of all human cancers. However, based on measurements reported three decades ago of Ras protein affinities to GTP in the 10-20 picomolar range, it has been accepted in the scientific and medical communities that Ras proteins are undruggable targets. Here, we report MicroScale Thermophoresis and scintillation proximity assay measurements of the affinity of K-Ras and several K-Ras mutants for GTP in the range of 200 nanomolar, a 10,000-fold difference from that previously reported, and the identification of over 400 small molecules that block GTP binding to K-Ras. Focusing on two of those molecules, we report small molecule inhibition of Ras downstream signaling and cellular proliferation in human pancreatic and non-small cell lung cancer cells expressing wild type and K-Ras G12C, G12D and G12S, and N-Ras Q61K mutants.

## INTRODUCTION

Ras proteins are encoded by three ubiquitously expressed genes (H-*RAS*, K-*RAS*, and N-*RAS*)(Parker and Mattos, 2018). They couple cell surface receptors to intracellular effector pathways and are key regulators of cellular growth and differentiation (Simanshu et al., 2017; Stephen et al., 2014). The binding of GTP and GDP, respectively, cycle Ras proteins between ‘on’ and ‘off’ signaling conformations. Under physiological conditions, the transition between these two states has been shown to be regulated by guanine nucleotide exchange factors (GEFs), which promote the activation of Ras proteins by stimulating the release of bound GDP and binding of GTP, and by GTPase-activating proteins (GAPs), which accelerate Ras-mediated GTP hydrolysis (Simanshu et al., 2017; Milburn et al., 1990; Shieh, 2019; Hunter et al., 2015). Ras proteins are a subset of the Ras GTPase superfamily which consists of over 150 human members which exhibit significant sequence homology in their GTP binding site, including Rac-1, Rho-A and cdc42 (Wennerberg et al., 2005; Cox and Der, 2010).

Approximately 30% of all human cancers are associated with mutations in Ras proteins (Prior et al., 2012; Hobbs et al., 2016; Pylayeva-Gupta et al., 2011). K-Ras is the most mutated Ras isoform. K-Ras mutations are frequently detected in pancreatic, colorectal and lung tumors (Prior et al., 2012; Hobbs et al., 2016; Cox et al., 2014). The most frequent K-Ras mutations are in residues Gly12, Gly13 and Gln61 (Hobbs et al., 2016; Li et al., 2018). N-Ras is mutated in about 20% of all melanoma patients (Jenkins and Sullivan, 2016). H-Ras mutations are comparatively rare (Prior et al., 2012).

A serious unmet medical need persists for patients with diseases, particularly cancer, associated with mutated *RAS* genes. A number of small molecules specifically targeting the K-Ras G12C mutant via a covalent mechanism have been identified, two of which are currently in clinical development (McCormick, 2020; Grapsa and Syrigos, 2020). However, potent small molecules targeting other mutants or wild-type Ras proteins have not been published. This is likely in major part due to the generally accepted paradigm that small molecule drugs cannot be developed which can compete with guanine nucleotides for binding to the GTP/GDP binding site of Ras, a conception based on studies from the 1990’s reporting GTP binding affinities for Ras proteins in the 10 – 20 pM range (John et al., 1990; John et al., 1993).

For the present study we used two contemporary technologies MicroScale Thermophoresis (MST) and scintillation proximity assay (SPA) to measure the affinity of K-Ras and several K-Ras mutants for GTP. Our measurements revealed dissociation constants (*K_d_*) for GTP binding that were 10^4^-fold higher than those reported previously, thus opening the way to target K-Ras proteins for drug development. Based on our findings, we developed a novel cell-free competitive binding assay to screen a focused small molecule library of 21,500 molecules and identified over 400 that compete with GTP binding to K-Ras, representing several different core structures. Based on their ability to compete with GTP binding to KRas, block K-Ras signaling, and inhibit cell proliferation in several tumor-derived human cell lines, we selected from these screening hits a subset for further SAR refinement. We report on the in vitro effects of two resulting molecules.

## RESULTS

### GTP affinity to wild-type K-Ras and three K-Ras mutants

Picomolar affinity data published in the 1990s for GTP binding to Ras were determined by filtration binding assays using [^3^H]-labeled nucleic acids (John et al., 1990; John et al., 1993). In the present study, we used MST and SPA to re-evaluate the *K_d_* of GTP binding to wild-type (WT) and three K-Ras mutants (G12D, G12C, Q61H) and two other Ras family members, Rac-1 and Rho-A. SPA is an established method that directly measures high-affinity ligand-protein interactions utilizing radiolabeled ligands and immobilized protein targets (Quick and Javitch, 2007; Rouck et al., 2017). It has been used extensively in drug discovery and high throughput screening of chemical libraries (Wu and Liu, 2005)(Glickman et al., 2008). MST is a validated, robust method for measuring K_d_’s in solution based on molecular motion induced by an applied temperature gradient (Jerabek-Willemsen et al., 2011). Several studies have compared affinity measurements by MST and Surface Plasmon Resonance (SPR) and found them to be identical (Wang et al., 2016); (Zhong et al., 2018)

Table 1 presents the affinities measured with the MST and SPA for K-Ras WT and three K-Ras mutants. In addition to the K-Ras WT, we studied the G12D, G12C, and Q61H mutants because they are the predominant forms of mutated K-Ras found in human tumors (Prior et al., 2012). The measured *K_d_* for wild type K-Ras by MST was approximately 460 nM and by SPA 250 nM. The measured *K_d_* values for the three tested K-Ras mutants ranged from approximately 120 nM to 280 nM by the two methods (Table 1). These values are orders of magnitude greater than the previously reported *K_d_* s. The measured *B_max_* values (Table 1) indicate a molar binding stoichiometry of GTP-to-target protein of unity, confirming that all tested recombinant proteins were purified in nucleic acid-free form.

**Table 1.** K_*d*_ and B_*max*_ values of GTP binding to several members of the Ras super family obtained by MST and SPA (nM)

| **Protein** | **MST** | **SPA** | **Bmax (mol:mol)** |
| --- | --- | --- | --- |
| K-Ras (wild type) | 463 ± 23 | 246 ± 19 | 1.00 ± 0.01 |
| K-Ras (G12D) | 244 ± 12 | 280 ± 19 | 0.97 ± 0.01 |
| K-Ras (G12C) | 207 ± 46 | 257 ± 20 | 0.94 ± 0.02 |
| K-Ras (Q61H) | 157 ± 21 | 118 ± 11 | 0.96 ± 0.02 |
| Rac-1 | 166 ± 10 | 151 ± 14 | 1.03 ± 0.02 |
| Rho-A | 130 ± 5 | 129 ± 12 | 1.02 ± 0.02 |

As a matter of reference, we also utilized MST and SPA to measure the affinities of Rac-1 and Rho-A for GTP. MST and SPA both yielded *K_d_* s consistent with previously published data for both GTPases (Zhang et al., 2000). As presented in Table 1, the *K_d_* for GTP binding by Rac-1 was approximately 170 nM by MST and 150 nM by SPA, while for Rho-A the *K_d_* value was approximately 130 nM by both methods. These data are comparable to those reported by Zhang et al (Zhang et al., 2000) using a filtration assay where Rac-1 and Rho-A exhibited *K_d_* s for GTP binding of 240 nM and 160 nM, respectively.

### Development of a novel GTP competition assay

The affinity data presented in Table 1 led us to conclude that the GTP binding site in members of the Ras superfamily could potentially serve as a target for small molecule drug development. We hypothesized that blocking GTP binding to Ras with a small molecule could render Ras in an inactive conformation, possibly similar to the GDP-bound conformation; block signaling along downstream pathways; and thereby exert a potential therapeutic benefit.

To identify small molecules that bind to the GTP binding site of wild type and mutant Ras proteins, we developed a cell-free competitive binding assay with which we screened a focused library of approximately 21,500 small molecules. The assay measured the ability of small molecules to compete with the binding of fluorescently (Cy3 or Cy5) labeled GTP to immobilized recombinant Ras (see Materials and Methods).

We miniaturized the assay to a 96-well format to support high-throughput screening and subsequent SAR efforts. Utilizing this competitive-binding assay, from the starting focused library we identified over 400 compounds that competed with GTP for binding to K-Ras WT and the K-Ras G12D, G12C and Q61H mutants. Structure analysis of these hits revealed 8 different core structures.

We subsequently developed compounds around the 8 core structures that robustly inhibit GTP binding to the K-Ras variants. Figure 1 and Table 2 summarize the *IC_50_* of two representative compounds (referred to as SHY-855 and SHY-867) that compete with GTP binding to K-Ras WT, the G12D, G12C and Q61H mutants, as well as Rac-1 and Rho-A, as measured by the competitive binding assay. Consistent with the inference drawn from the MST and SPA binding studies, the *IC_50_* values of these molecules demonstrate that small molecules can bind non-covalently to the GTP-binding site of Ras and competitively prevent GTP binding to that site. It should be noted that the measured *IC_50_* values in the sub μM range are a function of the optimized 100 nM concentration of the fluorescent-labeled GTP in the assay and not a measure of affinities.

**Fig 1.**
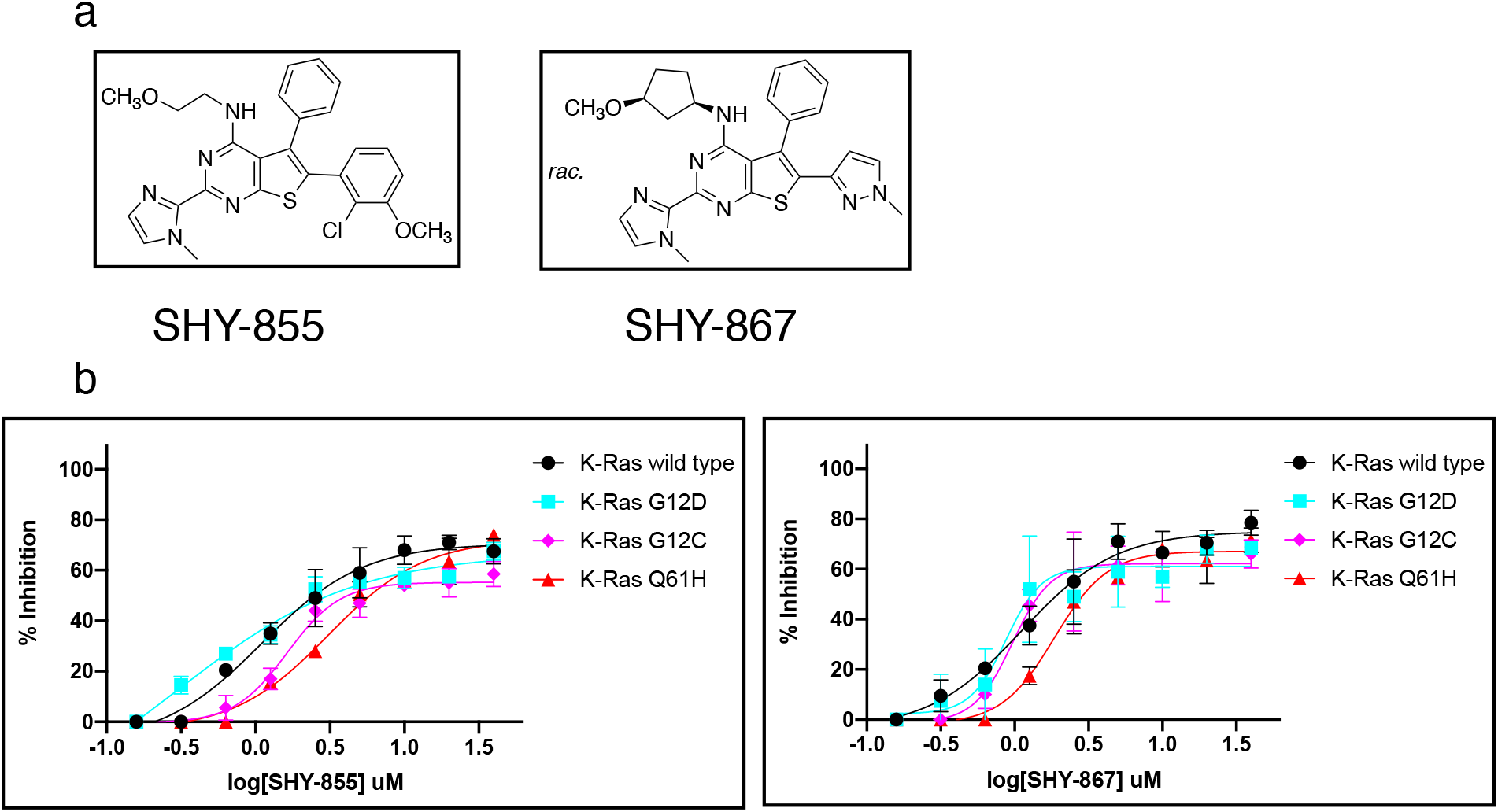
Two selected molecules compete for the GTP binding site of K-Ras proteins. a. Chemical structures of compounds SHY-855 and SHY-867. b. IC_*50*_ curves of the tested compounds blocking binding of fluorescent-labeled GTP to different K-Ras proteins.

**Table 2.** IC_*50*_ values of compounds competing with GTP (μM)

| **Protein** | **SHY-855** | **SHY-867** |
| --- | --- | --- |
| K-Ras (wild type) | 2.5 | 2.0 |
| K-Ras (G12D) | 2.5 | 2.5 |
| K-Ras (G12C) | 3.0 | 1.25 |
| K-Ras (Q61H) | 5.0 | 3.0 |
| Rac-1 | 2.5 | 2.0 |
| Rho-A | 2.5 | 2.5 |

Overall the two tested compounds SHY-855 and SHY-867 demonstrated similar *IC_50_* values, in the range of 1.25 – 5.0 μM, against the tested proteins (Fig. 1 and Table 2), suggesting they may be pan-Ras superfamily inhibitors. This may be explained by the high sequence homology of the GTP binding sites among these proteins.

### Inhibition of Ras downstream pathways

We have tested the inhibitory effects of compounds SHY-855 and SHY-867 on Ras-associated signal transduction pathways in three different human cell lines: the pancreatic cell-lines PANC-1 (K-Ras G12D mutant), MIA PaCa-2 (K-Ras G12C mutant), and the non-small cell lung cancer cell-line NCI-H1975 (KRas WT). Figures 2 and 3 illustrate the inhibitory effects of compounds SHY-855 and SHY-867 on phosphorylation and activation of MEK, Erk1/2 and Akt in those three cell lines. As shown in Figure 2, compound SHY-855 inhibited phosphorylation and activation of MEK, Erk1/2 and Akt in PANC-1, MIA PaCa-2 and NCI-H1975 cell lines in a dose-dependent manner with similar *IC_50_s* values (about 1-3 μM). These effects would be expected from a Ras inhibitor that is an upstream blocker of the two pathways and support the notion that compound SHY-855 induces an inactive conformation of Ras through a consistent binding mode, regardless of different mutations. In contrast, the *IC_50_s* of compound SHY-867 on the phosphorylation and activation of MEK and Erk1/2 were approximately 1 μM in PANC-1 and MIA PaCa-2 and 5 μM in NCI-H1975; and, of Akt, were approximately 7 μM in PANC-1 and MIA PaCa-2 and 10 μM in NCI-H1975 (Fig. 3). This data suggests a differential downstream signaling effect requiring further study and likely a more complicated explanation.

**Fig 2.**
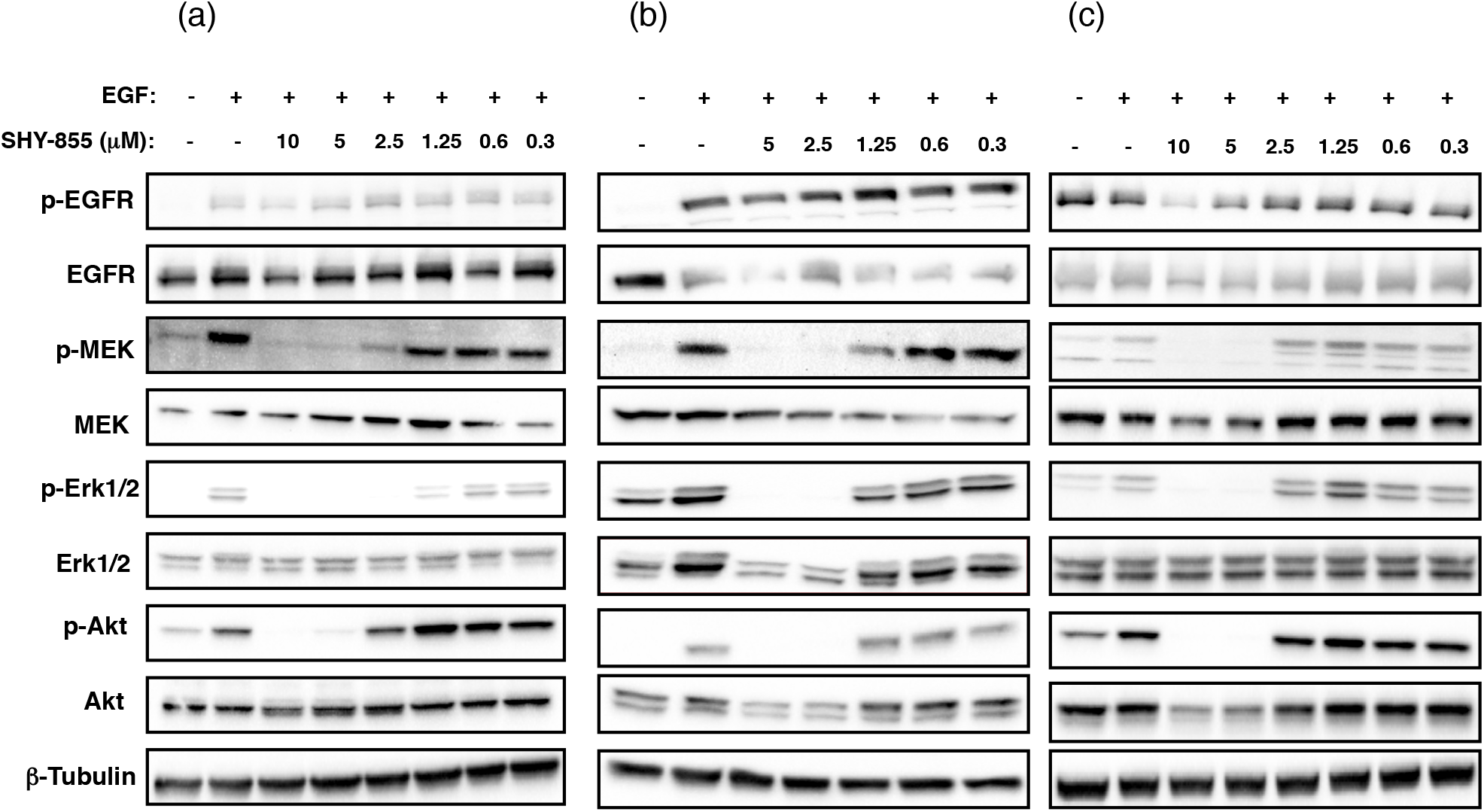
Compound SHY-855 inhibits K-Ras downstream signaling. Cells were incubated with compound SHY-855 for 6 hours with the indicated concentrations, stimulated with EGF for 15 minutes and cell lysates were analyzed using the indicated phospho-specific antibodies. (a) PANC-1 cell line, (b) MIA PaCa-2 cell line, (c) NCI-H1975 cell line.

**Fig 3.**
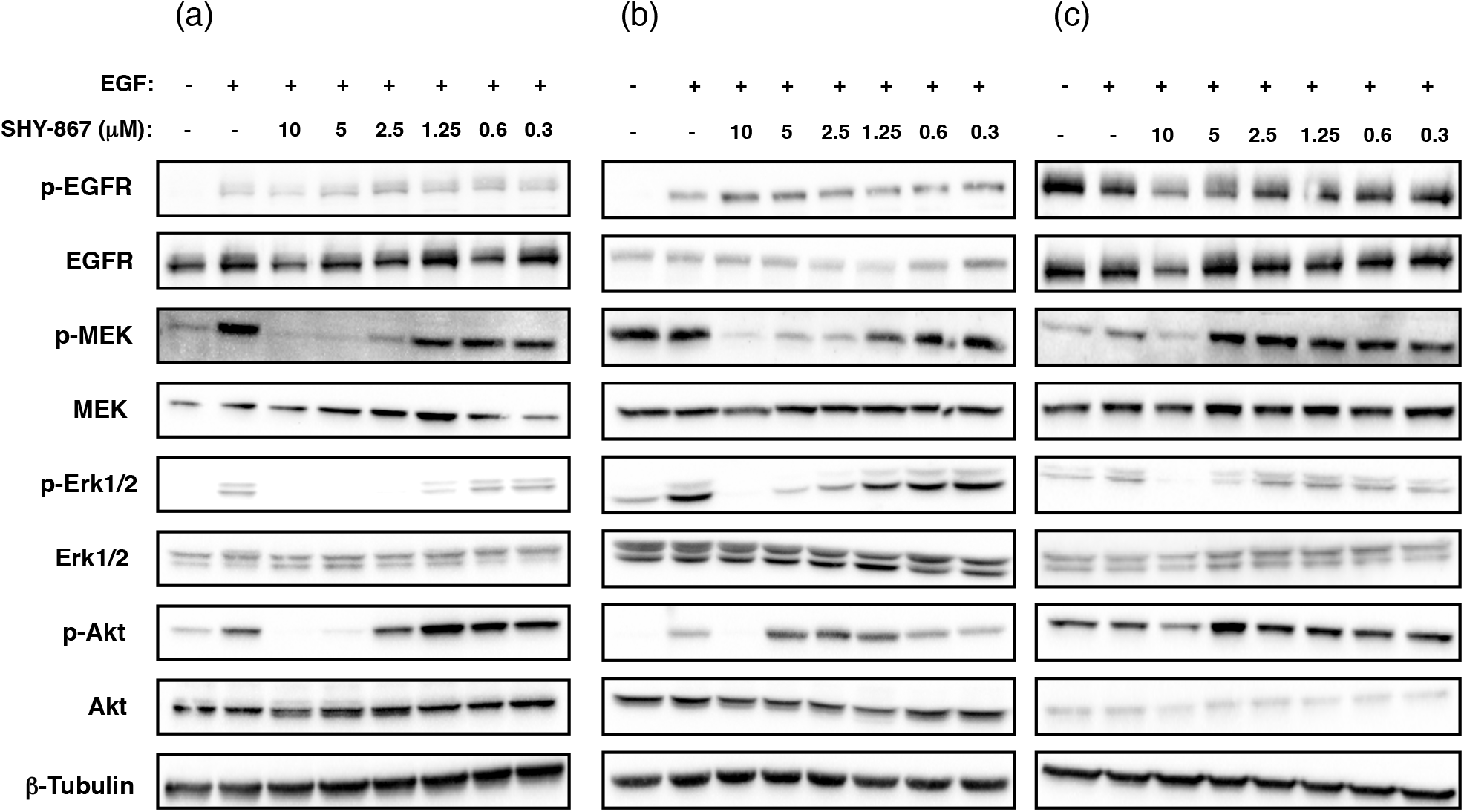
Compound SHY-867 inhibits K-Ras downstream signaling. Cells were incubated with compound SHY-867 for 6 hours with the indicated concentrations, stimulated with EGF for 15 minutes and cell lysates were analyzed using the indicated phosphospecific antibodies. (a) PANC-1 cell line, (b) MIA PaCa-2 cell line, (c) NCI-H1975 cell-line.

### Inhibition of the Ras-GTP complex

As a follow up to our downstream assays, we also measured the ability of compounds SHY-855 and SHY-867 to inhibit Ras-GTP complex formation in the three tested cell-lines (PANC-1, MIA PaCa-2 and NCI-H1975) using a Ras pull-down assay (see Material and Methods). As shown in Figure 4, both compounds prevent Ras-GTP complex formation with *IC_50_* values in the range of 2-3 μM. These values correlate well with the *IC_50_* values obtained for the inhibition of MEK, Erk1 /2 and Akt phosphorylation presented in Figures 2 and 3. The detected inhibition of Ras-GTP complex formation strongly suggests that both compounds bind directly to the GTP-binding site of Ras proteins and that, upon binding, the tested compounds induce a Ras inactive conformation, with a similar effect to the GDP-bound state.

**Fig 4.**
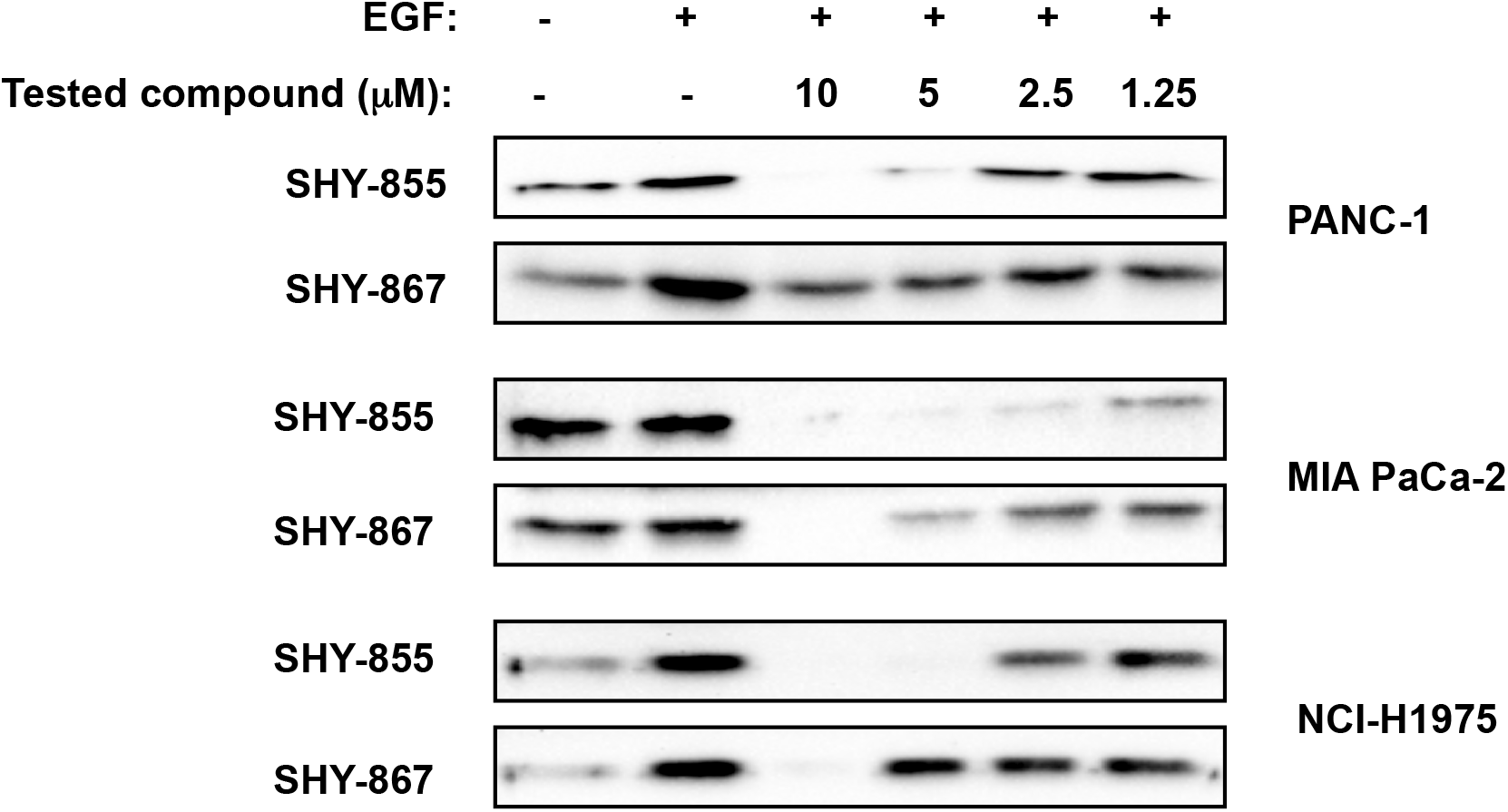
Compounds SHY-855 and SHY-867 inhibit Ras-GTP complex formation. The indicated cell lines were incubated with compound SHY-855 or SHY-867 for 6 hours with the indicated concentrations, and stimulated with EGF for 15 minutes. GTP pull down assay was performed on the cell lysate as described under Material and Methods.

### Anti-proliferative effects induced by the two selected compounds

We further measured the inhibition of compounds SHY-855 and SHY-867 on the cellular proliferation of six cell lines: the three cell lines mentioned above in the signaling studies (PANC-1, MIA PaCa-2, NCI-H1975), the pancreatic cell line BxPC3 (K-Ras WT) and the non-small cell lung cancer lines A549 (K-Ras G12S mutant) and NCI-H1299 (N-Ras Q61K mutant). Figure 5 and Table 3 summarize the *IC_50_* values measured in these studies. Molecules SHY-855 and SHY-867 both inhibit cellular proliferation in a dosedependent manner with similar *IC_50_* values, ranging from approximately 0.45 μM to 1.20 μM in all tested cell lines. Consistent with the notion that these two compounds are pan-Ras superfamily inhibitors, it is reasonable to speculate that they can inhibit pathways downstream of several members of the Ras superfamily. Interestingly, in contrast to the different *IC_50_* values obtained for inhibition of phosphorylation and activation of MEK, Erk1/2 and Akt presented in Table 2, the *IC_50_* values for inhibition of proliferation are more similar.

**Fig 5.**
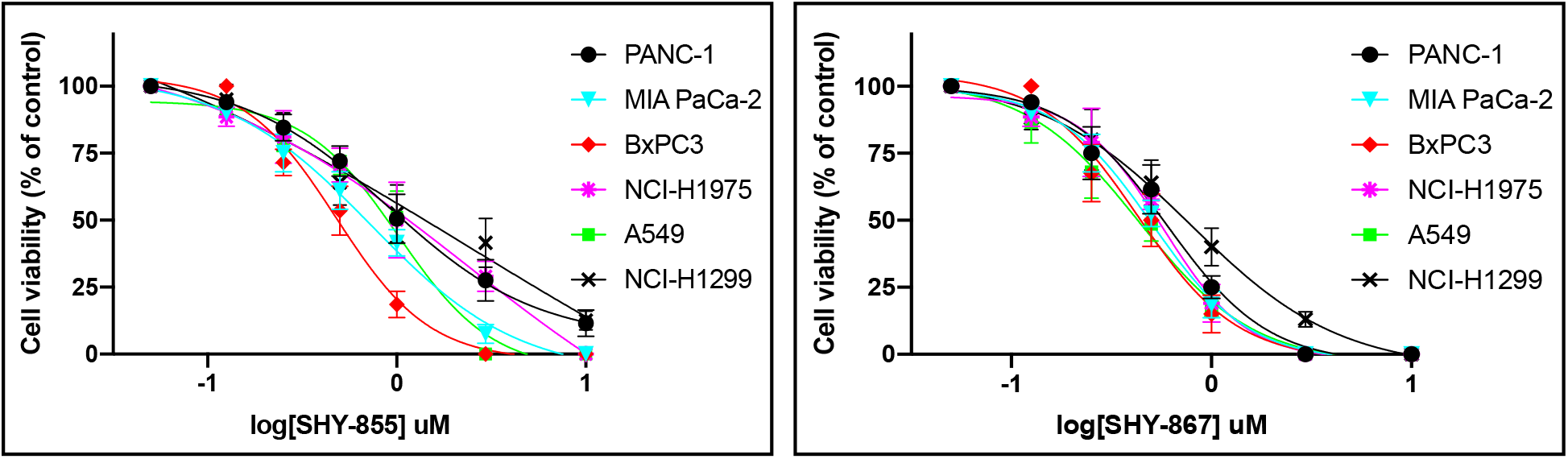
Effects of compounds SHY-855 and SHY-867 on cellular viability. The indicated cell lines were incubated with compound SHY-855 (a) or SHY-867 (b) for 72 hours with the indicated concentrations. Following 72 hours of incubation cell viability was analyzed as described under Material and Methods.

**Table 3.** IC_*50*_ values of proliferation assays (nM)

|  | **PANC-1** | **MIA PaCa-2** | **BxPC-3** | **NCI-H1975** | **A549** | **NCI-H1299** |
| --- | --- | --- | --- | --- | --- | --- |
| **SHY-855** | **1000** | **750** | **600** | **1000** | **1000** | **1200** |
| **SHY-867** | **750** | **650** | **550** | **600** | **450** | **800** |

## DISCUSSION

The data presented here demonstrates for the first time that small molecules can be found that compete with GTP for binding to the GTP binding site of wild type and several K-Ras mutants.

Using small molecules to compete for the ATP binding site of kinases is a well-validated drug development approach which has yielded a significant number of FDA approved drugs (Wu et al., 2016; Roskoski, 2020). Unlike kinases, where displacement of ATP by a small molecule is sufficient to block phosphorylation and activation events, to block Ras activation, antagonists targeting the GTP binding site of Ras must both bind and induce or stabilize an inactive conformation with similar effect to the GDP-bound inactive conformation of the GTPase. Since blocking GTP binding in our cell-free assay cannot predict which conformation is induced by a compound, inhibitory activity also had to be assessed in cell-based assays. The demonstrated cell-based activity including inhibition of MEK, Erk1/2 and Akt phosphorylation, Ras-GTP complex formation and proliferation reported here, demonstrate that the tested small molecules can penetrate tumor cells and induce or stabilize an inactive Ras conformation. Analysis of the measured *IC_50_* values of the two tested small molecules in the different assays indicates that they are most active in the proliferation assays. In both cell-free competition assays and cell-based phosphorylation assays, the measured *IC_50_* values are in the sub μM range, while in the proliferation assays they are in the high nM range. These results may reflect the pan-Ras superfamily inhibitory effects of the molecules tested.

Recently reported small molecules targeting the K-Ras G12C mutant (Ostrem et al., 2013; McCormick, 2020; Canon et al., 2019; Hallin et al., 2020; Lito et al., 2016) are different from the molecules described here. They are specific to only one mutant, bind covalently, are not competitive binders, and lock the protein in an inactive conformation with GDP bound.

While further work remains to be done, this paper strongly suggests that opportunities should exist to develop small molecule therapeutics that address clear unmet medical needs, including cancers and inflammatory diseases, by directly targeting the GTP binding site of K-Ras and other small GTPases that are members of the Ras superfamily.

## Materials and Methods

### Reagents

Human tumor-derived NSCLC cells NCI-H1975 (RRID:CVCL_UE30), A549 (RRID:CVCL_0023), NCI-H1299 (RRID:CVCL_0060) and pancreatic cancer cell lines PANC-1 (RRID:CVCL_0480), MIA-PaCa-2 (RRID:CVCL_HA89) and BxPC3 (RRID:CVCL_0186), were all purchased from American Type Culture Collection and grown in complete RPMI medium (NCI-H1975, NCI-H1299, BxPC3), DMEM-High Glucose (PANC-1 and MIA-PaCa-2) or F-12K medium (A549), supplemented with penicillin (100 U/mL), streptomycin (100 μg/ml), and 10% heat-inactivated FBS at 37 °C in a humidified incubator with 5 % CO_2_.

The following antibodies were used: rabbit mAb anti-human phospho-EGF Receptor (Tyr1068)(RRID:AB_2096270); rabbit mAb anti-human EGFR (RRID:AB_2246311); rabbit mAb antihuman phospho-AKT (Ser473)(RRID:AB_2315049); rabbit mAb anti-human pan-AKT (RRID:AB_2225340); rabbit mAb anti-human phospho-p44/42 ERK (Thr202/Tyr204) (RRID:AB_331772); rabbit Ab anti-human p44/42 ERK (RRID:AB_330744); rabbit Ab anti-human phospho-MEK1/2 (Ser217/221) (RRID:AB_331648), rabbit mAb anti-human MEK1/2 (RRID:AB_331778); mouse mAb anti-human RAS (Cell Signaling Technology). Compounds SHY-855 and SHY-867 were synthesized at Enamine LLC (Kiev, Ukraine) and were over 95% pure.

Proteins expression: K-Ras wild type and mutants (residues 2-185), Rac1 (residues 1-177), RhoA (residues 2-181), were subcloned into the MCSG7 bacterial expression vector which encodes an N-terminal deca-histidine fusion tag followed by a TEV protease cleavage site. All proteins were expressed in *E. coli* BL21 (DE3) gold cells, purified by nickel-affinity and size-exclusion chromatography on an AKTAxpress system (GE Life Sciences), which consisted of a 1mL nickel affinity column followed by a Superdex 200 16/60 gel filtration column. The final protein buffer consisted of 50 mM Tris-HCl pH 7.5, 150 mM NaCl, 1 mM MgCl_2_ and 1 mM DTT (Buffer I).

### Kinetic studies

MicroScale Thermophoresis experiments were performed according to the NanoTemper Technologies protocol in a Monolith NT.115 (red) instrument (NanoTemper Technologies, Munich, Germany). Binding was measured between EDA-GTP-Cy5 (Jena Bioscience GmbH, Jena, Germany) and His-tagged labeled proteins in Buffer I. Data analyses and the dissociation constants (*K_d_*) calculation were performed using MO Affinity Analysis software (NanoTemper Technologies, Munich, Germany). Binding of radiolabeled GTP was measured by means of the SPA. 25 ng of purified recombinant protein were immobilized on 1.25 mg copper-coated His tag SPA beads per mL of assay buffer (20 mM Tris-Cl, pH 7.45, 150 mM NaCl, 3 mM MgCl_2_, 1 mM TCEP) in 100 μL-assays. In pilot experiments this amount of protein was determined to prevent radioligand depletion and yield a level of protein occupation far below the binding capacity of the beads. YSi (Yttrium silicate) copper-coated His-tag beads (Perkin Elmer) were used in combination with α[^32^P]GTP (American Radiolabeled Chemicals, Inc.; 3000 Ci/mmol) and PVT (polyvinyl toluene). All binding assays were performed in 96-well white wall clear-bottom plates and assayed in a Wallac photomultiplier tube MicroBeta™ counter. Saturation binding experiments were performed with increasing concentrations of radiolabeled GTP (2.5 nM – 10 μM), and nonspecific binding (the non-proximity signal) was assayed in the presence of 800 mM imidazole. The stoichiometry of radiolabeled GTP binding to the GTPases was determined by using known amounts of protein (1.2 pmol) and calculating the specific binding (total signal – non-proximity signal) at each tested GTP concentration. Counts-per-minute (cpm) were transformed into pmol using known concentrations of radiolabeled GTP to determine the efficiency of detection. Non-linear regression fitting in GraphPad Prism 7 was used to calculate the dissociation constant (*K_d_*) and maximum binding (*B_max_*) by plotting the bound pmol of GTP as a function of free GTP. Data originate from three independent experiments that were performed with technical triplicates.

### Cell-free assay for GTP-binding proteins

Tested purified proteins were all His-tagged and diluted to 3-10 μg/ml solution in Buffer I containing 50 mM Tris, pH 7.5, 150 mM NaCl, 1 mM MgCl2 and 1 mM DTT. 200 μl of the diluted His-tag protein was then added to a nickel coated 96 well plate and incubated overnight at 4 °C. The next day, the wells were triple washed in 200 μl of Buffer I and then 200 μl of Buffer I were added to each well in the presence of 1% DMSO. The tested molecules were added to the protein-coated wells at final concentration of 20 μM. For IC50 measurements serial dilutions of all tested small molecules were prepared and added to the protein-coated wells. Following 3 hours of incubation at room temperature, Cy3-GTP or Cy5-GTP was added to each well in a final concentration of 100 nM. The labeled GTP was incubated for 45 min at 23 °C. Then, wells were washed 3X in Buffer I, and 200 μl of Buffer I were added to each well and the amount of bound labeled-GTP was measured with a Molecular Devices SpectraMax M3 plate reader.

### Cell-based phosphorylation assays

Cells were plated at 350000 cells/well density in a 12-well plate, allowed 3 hours to adhere to the plate, then starved in the appropriate medium in the presence of 0.5% FBS overnight. Serial dilutions of small molecules to be tested were added to cells in the presence of 0.3% DMSO for 6 hours incubation at 37 °C. Next, cells were stimulated with 1.5 ng/ml EGF for 15 minutes then cells were lysed with lysis buffer containing 1% Triton X-100, EDTA, and Halt™ Protease & Phosphatase Inhibitor Cocktail (Thermo Scientific). Protein concentration was assessed by BCA protein assay (Thermo Scientific).

### Western Blotting

Equal volumes of eluate (25 μl) were separated by 10 % SDS-PAGE and transferred to nitrocellulose membranes (Invitrogen by Thermo Fisher Scientific). The membrane was stained with Ponceau S Stain (Boston BioProducts) to verify uniform protein loading. Membranes were blocked with 5 % BSA in TBST and then incubated overnight at 4 °C with primary antibody, followed by HRP-conjugated secondary antibody (Jackson Immunoresearch, West Grove, PA). Membranes were incubated in Amersham ECL Prime Western Blotting Detection Reagent (GE Healthcare) and bands were visualized using the ChemiDoc MP imaging system (Bio-Rad).

### GTP pull down assay

Cells were plated at 2•10^6^ cells/well density in a 6-well plate, allowed 3 hours to adhere to the plate, then starved in the appropriate medium in the presence of 0.5 % FBS overnight. Serial dilutions of the small molecules to be tested were added to the cells in the presence of 0.3% DMSO for 6 hours incubation at 37 °C. Next, cells were stimulated with 5 ng/ml EGF for 15 minutes, rinsed with ice-cold PBS and then lysed with 500 μl of lysis/binding/wash buffer (25 mM Tris-HCl, pH 7.2, 150 mM NaCl, 5 mM MgCl2, 5 % glycerol, 1 % NP40) from Active Ras Detection kit (Cell Signaling Technology) supplemented with Halt™ Protease & Phosphatase Inhibitor Cocktail (Thermo Scientific). To account for significant differences in cell number due to the treatment, a small sample of lysate was saved for protein quantification and the rest of the lysate was snap frozen. Protein concentration was assessed by BCA protein assay (Thermo Scientific). To ensure that equal amount of protein undergoes RBD pulldown, lysates were subsequently thawed (at 23 °C) and adjusted to 1 mg/ml with lysis/binding/wash buffer (0.5 ml volume). Equal amounts of lysate were then added to 0.5 mL lysis buffer containing RAF-RBD (1 mL total volume). Lysates were vortexed, incubated for 10 min on ice and subsequently pre-cleared at 14,000 rpm for 5 min at 4 °C. 90% of the pre-cleared lysates were subsequently added to prewashed glutathione agarose beads from Active Ras Detection kit (Cell signaling Technology, #8821) for 1 hour at 4 °C under constant rocking. The beads were subsequently pelleted, washed 3 times with lysis/binding/wash buffer, and eluted for western blotting with 50 μl of 1X SDS-PAGE sample buffer. Level of GTP-bound RAS was determined by western blot.

### Proliferation assay

Cells were plated at 4000 cells/well density in 96-wells plate. The next day serial dilutions of the tested small molecules were added to the cells in the presence of 0.3 % DMSO and 10 % FBS. After small molecule addition, cells were incubated for 3 days at 37 °C in a humidified incubator with 5 % CO_2_. At the end of the incubation period, cell viability was measured using the CellTiter 96^®^ Aqueous One Solution Cell Proliferation Assay according to manufacturer specifications (Promega, Madison, WI).

## ACKNOWLEGMENT

We thank Allan S. Jacobson (University of Massachusetts, Worcester) for critical comments on the manuscript.

## References

Canon, J., Rex, K., Saiki, A.Y., Mohr, C., Cooke, K., Bagal, D., Gaida, K., Holt, T., Knutson, C.G., Koppada, N., Lanman, B.A., Werner, J., Rapaport, A.S., San Miguel, T., Ortiz, R., Osgood, T., Sun, J.-R., Zhu, X., McCarter, J.D., Volak, L.P., Houk, B.E., Fakih, M.G., O’Neil, B.H., Price, T.J., Falchook, G.S., Desai, J., Kuo, J., Govindan, R., Hong, D.S., Ouyang, W., Henary, H., Arvedson, T., Cee, V.J., Lipford, J.R., 2019. The clinical KRAS(G12C) inhibitor AMG 510 drives anti-tumour immunity. Nature 575, 217–223. https://doi.org/10.1038/s41586-019-1694-1

Cox, A.D., Der, C.J., 2010. Ras history: The saga continues. Small GTPases 1, 2–27. https://doi.org/10.4161/sgtp.1.1.12178

Cox, A.D., Fesik, S.W., Kimmelman, A.C., Luo, J., Der, C.J., 2014. Drugging the undruggable RAS: Mission possible? Nat. Rev. Drug Discov. 13, 828–851. https://doi.org/10.1038/nrd4389

Glickman, J.F., Schmid, A., Ferrand, S., 2008. Scintillation proximity assays in high-throughput screening. Assay Drug Dev. Technol. 6, 433–455. https://doi.org/10.1089/adt.2008.135

Grapsa, D., Syrigos, K., 2020. Direct KRAS inhibition: progress, challenges and a glimpse into the future. Expert Rev. Anticancer Ther. https://doi.org/10.1080/14737140.2020.1760093

Hallin, J., Engstrom, L.D., Hargis, L., Calinisan, A., Aranda, R., Briere, D.M., Sudhakar, N., Bowcut, V., Baer, B.R., Ballard, J.A., Burkard, M.R., Fell, J.B., Fischer, J.P., Vigers, G.P., Xue, Y., Gatto, S., Fernandez-Banet, J., Pavlicek, A., Velastagui, K., Chao, R.C., Barton, J., Pierobon, M., Baldelli, E., Patricoin, E.F., Cassidy, D.P., Marx, M.A., Rybkin, I.I., Johnson, M.L., Ou, S.-H.I., Lito, P., Papadopoulos, K.P., Jänne, P.A., Olson, P., Christensen, J.G., 2020. The KRASG12C Inhibitor MRTX849 Provides Insight toward Therapeutic Susceptibility of KRAS-Mutant Cancers in Mouse Models and Patients. Cancer Discov. 10, 54–71. https://doi.org/10.1158/2159-8290.CD-19-1167

Hobbs, G.A., Der, C.J., Rossman, K.L., 2016. RAS isoforms and mutations in cancer at a glance. J. Cell Sci. 129, 1287–1292. https://doi.org/10.1242/jcs.182873

Hunter, J.C., Manandhar, A., Carrasco, M.A., Gurbani, D., Gondi, S., Westover, K.D., 2015. Biochemical and Structural Analysis of Common Cancer-Associated KRAS Mutations. Mol. Cancer Res. MCR 13, 1325–1335. https://doi.org/10.1158/1541-7786.MCR-15-0203

Jenkins, R.W., Sullivan, R.J., 2016. NRAS mutant melanoma: an overview for the clinician for melanoma management. Melanoma Manag. 3, 47–59. https://doi.org/10.2217/mmt.15.40

Jerabek-Willemsen, M., Wienken, C.J., Braun, D., Baaske, P., Duhr, S., 2011. Molecular interaction studies using microscale thermophoresis. Assay Drug Dev. Technol. 9, 342–353. https://doi.org/10.1089/adt.2011.0380

John, J., Rensland, H., Schlichting, I., Vetter, I., Borasio, G.D., Goody, R.S., Wittinghofer, A., 1993. Kinetic and structural analysis of the Mg(2+)-binding site of the guanine nucleotide-binding protein p21H-ras. J. Biol. Chem. 268, 923–929.

John, J., Sohmen, R., Feuerstein, J., Linke, R., Wittinghofer, A., Goody, R.S., 1990. Kinetics of interaction of nucleotides with nucleotide-free H-ras p21. Biochemistry 29, 6058–6065. https://doi.org/10.1021/bi00477a025

Li, S., Balmain, A., Counter, C.M., 2018. A model for RAS mutation patterns in cancers: finding the sweet spot. Nat. Rev. Cancer 18, 767–777. https://doi.org/10.1038/s41568-018-0076-6

Lito, P., Solomon, M., Li, L.-S., Hansen, R., Rosen, N., 2016. Allele-specific inhibitors inactivate mutant KRAS G12C by a trapping mechanism. Science 351, 604–608. https://doi.org/10.1126/science.aad6204

McCormick, F., 2020. Sticking it to KRAS: Covalent Inhibitors Enter the Clinic. Cancer Cell 37, 3–4. https://doi.org/10.1016/j.ccell.2019.12.009

Milburn, M.V., Tong, L., deVos, A.M., Brünger, A., Yamaizumi, Z., Nishimura, S., Kim, S.H., 1990. Molecular switch for signal transduction: structural differences between active and inactive forms of protooncogenic ras proteins. Science 247, 939–945. https://doi.org/10.1126/science.2406906

Ostrem, J.M., Peters, U., Sos, M.L., Wells, J.A., Shokat, K.M., 2013. K-Ras(G12C) inhibitors allosterically control GTP affinity and effector interactions. Nature 503, 548–551. https://doi.org/10.1038/nature12796

Parker, J.A., Mattos, C., 2018. The K-Ras, N-Ras, and H-Ras Isoforms: Unique Conformational Preferences and Implications for Targeting Oncogenic Mutants. Cold Spring Harb. Perspect. Med. 8. https://doi.org/10.1101/cshperspect.a031427

Prior, I.A., Lewis, P.D., Mattos, C., 2012. A comprehensive survey of Ras mutations in cancer. Cancer Res. 72, 2457–2467. https://doi.org/10.1158/0008-5472.CAN-11-2612

Pylayeva-Gupta, Y., Grabocka, E., Bar-Sagi, D., 2011. RAS oncogenes: weaving a tumorigenic web. Nat. Rev. Cancer 11, 761–774. https://doi.org/10.1038/nrc3106

Quick, M., Javitch, J.A., 2007. Monitoring the function of membrane transport proteins in detergent-solubilized form. Proc. Natl. Acad. Sci. U. S. A. 104, 3603–3608. https://doi.org/10.1073/pnas.0609573104

Roskoski, R., 2020. Properties of FDA-approved small molecule protein kinase inhibitors: A 2020 update. Pharmacol. Res. 152, 104609. https://doi.org/10.1016/j.phrs.2019.104609

Rouck, J.E., Krapf, J.E., Roy, J., Huff, H.C., Das, A., 2017. Recent advances in nanodisc technology for membrane protein studies (2012-2017). FEBS Lett. 591, 2057–2088. https://doi.org/10.1002/1873-3468.12706

Shieh, J.T.C., 2019. Emerging RAS superfamily conditions involving GTPase function. PLoS Genet. 15, e1007870. https://doi.org/10.1371/journal.pgen.1007870

Simanshu, D.K., Nissley, D.V., McCormick, F., 2017. RAS Proteins and Their Regulators in Human Disease. Cell 170, 17–33. https://doi.org/10.1016/j.cell.2017.06.009

Stephen, A.G., Esposito, D., Bagni, R.K., McCormick, F., 2014. Dragging ras back in the ring. Cancer Cell 25, 272–281. https://doi.org/10.1016/j.ccr.2014.02.017

Wang, X., Wu, X., Zhang, A., Wang, S., Hu, C., Chen, W., Shen, Y., Tan, R., Sun, Y., Xu, Q., 2016. Targeting the PDGF-B/PDGFR-β Interface with Destruxin A5 to Selectively Block PDGF-BB/PDGFR-ββ Signaling and Attenuate Liver Fibrosis. EBioMedicine 7, 146–156. https://doi.org/10.1016/j.ebiom.2016.03.042

Wennerberg, K., Rossman, K.L., Der, C.J., 2005. The Ras superfamily at a glance. J. Cell Sci. 118, 843–846. https://doi.org/10.1242/jcs.01660

Wu, P., Nielsen, T.E., Clausen, M.H., 2016. Small-molecule kinase inhibitors: an analysis of FDA-approved drugs. Drug Discov. Today 21, 5–10. https://doi.org/10.1016/j.drudis.2015.07.008

Wu, S., Liu, B., 2005. Application of scintillation proximity assay in drug discovery. BioDrugs Clin. Immunother. Biopharm. Gene Ther. 19, 383–392. https://doi.org/10.2165/00063030-200519060-00005

Zhang, B., Zhang, Y., Wang, Z., Zheng, Y., 2000. The role of Mg2+ cofactor in the guanine nucleotide exchange and GTP hydrolysis reactions of Rho family GTP-binding proteins. J. Biol. Chem. 275, 25299–25307. https://doi.org/10.1074/jbc.M001027200

Zhong, W., Sun, B., Gao, W., Qin, Y., Zhang, H., Huai, L., Tang, Y., Liang, Y., He, L., Zhang, X., Tao, H., Chen, S., Yang, W., Yang, L., Liu, Y., Liu, H., Zhou, H., Sun, T., Yang, C., 2018. Salvianolic acid A targeting the transgelin-actin complex to enhance vasoconstriction. EBioMedicine 37, 246–258. https://doi.org/10.1016/j.ebiom.2018.10.041

